# Evolutionary transition of *doublesex* regulation in termites and cockroaches: from sex-specific splicing to male-specific transcription

**DOI:** 10.1101/2021.05.31.446247

**Authors:** Satoshi Miyazaki, Kokuto Fujiwara, Keima Kai, Yudai Masuoka, Hiroki Gotoh, Teruyuki Niimi, Yoshinobu Hayashi, Shuji Shigenobu, Kiyoto Maekawa

## Abstract

The sex determination gene *doublesex* (*dsx*), which encodes a transcription factor with two domains: oligomerisation domain 1 (OD1) and OD2, is conserved among insects. The sex-specific Dsx splicing isoforms regulate the transcription of target genes and trigger sex differentiation in all holometabolous insects examined to date. However, in some hemimetabolous insects, *dsx* is less conserved and not spliced sexually. Here, to elucidate evolutionary changes in *dsx* in the gene structure and its regulatory manner in termites, we searched genome and/or transcriptome databases for the OD1 and OD2 of *dsx* in six termite species and their sister group (woodroach). Molecular phylogenetic analysis identified OD1 sequences of termites and a woodroach clustered with *dsx* of holometabolous insects and regarded them as *dsx* orthologues. In the woodroach, *a dsx* orthologue containing OD2 was spliced sexually, as previously shown in other insects. However, OD2 were not found in all termite *dsx* orthologues. These orthologues were encoded by only a single exon in three termites with genome information; they were not alternatively spliced, but transcribed in a male-specific manner in two species examined. Evolution of *dsx* regulation from sex-specific splicing to male-specific transcription might be occurred at the early stage of social evolution in termites.

## Introduction

In insects, sex determination is cell-autonomously controlled by a cascade composed of several genes, where the *doublesex* (*dsx*) encoding sex-specific transcription factors is located at the bottom ^1^. In the cascade of fruit fly (*Drosophila melanogaster*), the primary signal, which is the number of X chromosome ^2^ is sequentially transduced into the sex-specific splicing of *Sex-lethal, transformer*, and *dsx*, resulting in the sex-specific transcription of target genes responsible for their sex differentiation ^3,4^. Although upstream genes encoding splicing regulators in this cascade differ among insect taxa, *dsx* is conserved among them, as well as in some crustaceans and chelicerates that are non-insect arthropods ^1,5–7^. Moreover, the *dsx* orthologue is sexually spliced in all holometabolous insects examined to date ^3,4^, thus, such *dsx* regulation is believed to be a general feature of the cascade in all insects. However, in hemimetabolous insects, the phylogenetic position of which is the interval between holometabolous insects and non-insect arthropods, *dsx* is sexually spliced in two of four species examined recently, but spliced alternatively and not sex-specifically in the others ^8,9^. Furthermore, in some crustaceans and mites, *dsx* does not produce sex-specific splicing isoforms and is instead expressed in a sex-specific manner, which controls their male differentiation ^5,7^. Therefore, in contrast to the knowledge based on holometabolous insects, the sex-specific regulation of *dsx* would be evolutionarily labile among insects and non-insect arthropods, and it could be diversified in hemimetabolous insects.

Price et al. ^6^ searched public databases of insect genomes and transcriptomes for *dsx* homologues, based on the presence of two domains conserved throughout holometabolous insects: DNA-binding DM domain and dimerisation domain, referred to as oligomerisation domain (OD) 1 and OD2, respectively. OD1 is shared among not only *dsx* but also *doublesex/mab-3 related transcription factor* (*DMRT*) genes that are conserved among metazoans and involved in their sexual development ^10,11^, whereas OD2 is specific to *dsx* ^12^. They failed to obtain any evidence of the presence of *dsx* in several hemimetabolous insects, including termites (infraorder Isoptera or epifamily Termitoidae) belonging to the monophyletic group within the cockroach clade ^13,14^. In the lower (primitive) termite *Zootermopsis nevadensis*, neither domain was found, and only OD1 was found in the higher (derived) termite *Nastitermes takasagoensis* ^6^. The genome sequence of the German cockroach *Blattella germanica* has been published, and a *dsx* homologue has been identified in the genome (gene ID: PSN43312.1 ^15^). *Blattella dsx* was shown to contain both OD1 and OD2 (referred to as “DM domain” and “Dsx Dimerization domain”, respectively, in Wexler et al. ^9^) and to be sexually spliced as in holometabolous insects ^9^. Based on these results, we hypothesised that the *dsx* sequence (especially OD1 and/or OD2) diversified during termite evolution, and then *dsx* sequence diversification affected its regulatory manner. These hypotheses could be examined by a more comprehensive search of complete genome sequences and transcriptomic data in termites and in their sister group, woodroach (*Cryptocercus* spp.) ^16–21^.

The present study aimed to elucidate evolutionary changes in *dsx* in the gene structure and its regulatory manner in the clade of cockroaches and termites. First, we exhaustively searched genome and/or transcriptome databases of the woodroach *Cryptocercus punctulatus* and six termite species (*Reticulitermes speratus*, *Hodotermopsis sjostedti*, *Cryptotermes secundus*, and *Macrotermes natalensis*, as well as *Z. nevadensis* and *N. takasagoensis*) for the OD1 and OD2 of *dsx*, and then examined whether the searched sequences were *dsx* homologues based on the molecular phylogeny and synteny. Second, we performed gene expression analysis based on quantitative RT-PCR and published transcriptome data, for one woodroach species and two termite species. Third, to identify putative regulatory factors in the sex-specific transcription of termite *dsx*, transcription factor binding sites (TFBSs) were identified in the promoter region of the *dsx* orthologue of *R. speratus*. The expression levels of putative transcription factors were compared between the sexes based on published transcriptome data. On the basis of the results obtained, we concluded that *dsx* regulation shifted from sex-specific splicing to male-specific transcription at the early stage of social evolution in termites.

## Results

### Searches for *doublesex* orthologues in termites and woodroach

We performed BLAST searches using the translated OD1 sequence of *Blattella dsx* (45 amino acids) as a query against genome and/or transcriptome databases of six termite species and one woodroach species, after which two to four OD1 sequences were hit in each species. Then, BLAST searches were performed using the translated OD2 sequence of *Blattella dsx* (45 amino acids) as a query. A single sequence from the woodroach was hit (Cpun_comp8195_c0_seq1, Table 1), but no sequence was hit from any termites. Rapid amplification cDNA ends (RACE) PCR was performed using primers specific for Cpun_comp8195_c0_seq1 (Table S1), and a single full-length transcript was obtained only from females. Then, using a reverse primer specific to the male-specific exon of *Blattella dsx* (located at terminal codon and 3’UTR, Table S1), another 3’ end of coding sequence was amplified from males. As these transcripts contained an OD1 upstream of OD2 (Fig. 1), it was designated as *C. punctulatus dsx* (*Cpun_dsx*). The amplified sequences downstream of OD2 were predicted to be the sex-specific exon(s) of *Cpun_dsx*. The determined nucleotide and putative amino acid sequences of the *Cpun_dsx* splicing variants are available in the DDBJ/EMBL/GenBank databases (accession no. xxxxx and xxxxxx).

**Figure 1.**
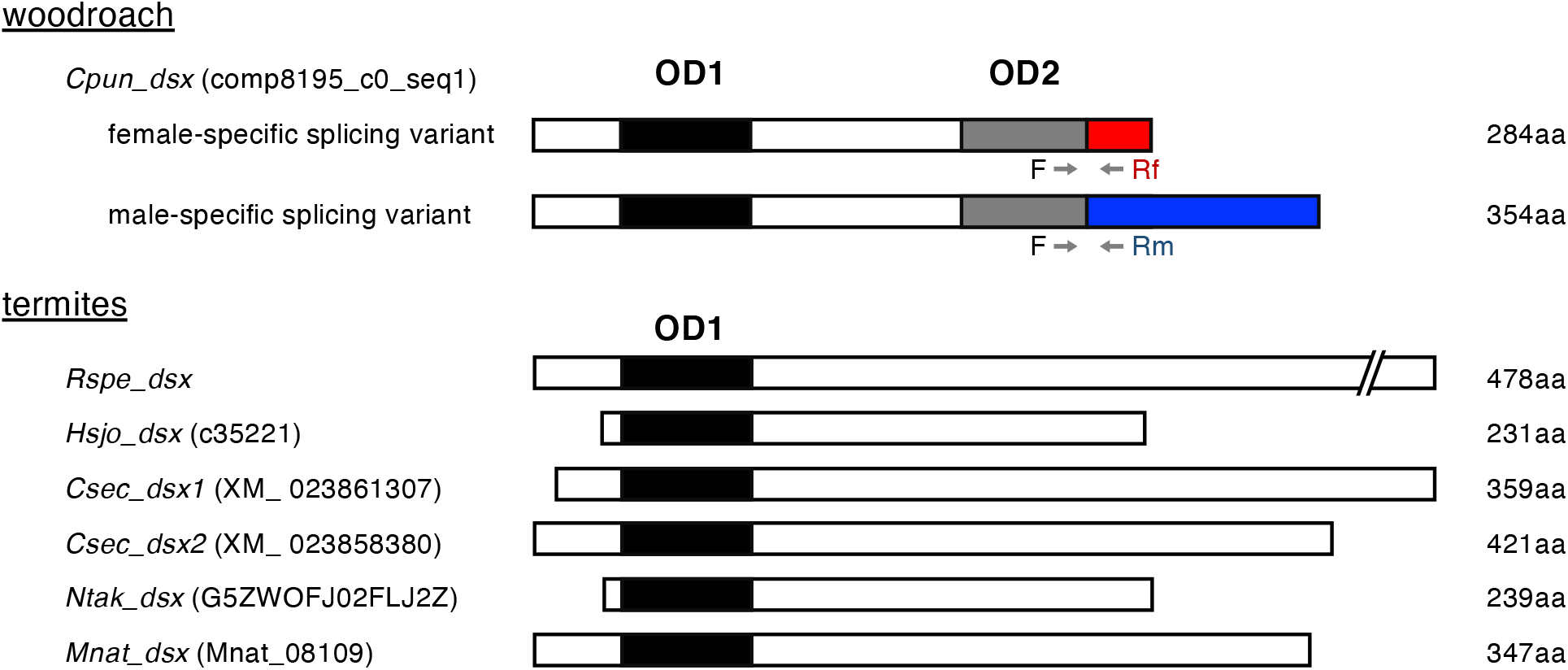
Structures of *dsx* transcripts in woodroach and termites. Full-length coding sequence of *dsx* transcripts in one woodroach and five termite species are shown as transcribed from left to right. The OD1 is shown as a filled box, whereas the sexually shared, female-specific, and male-specific OD2 is shown as a grey, red, and blue box, respectively. Numbers on the right of boxes are amino acid lengths of the translated *dsx*. Positions of primers for quantifying *Cpun_dsx* expression by quantitative RT-PCR are shown in arrows: “F”, “Rf”, and “Rm” indicate the shared forward primer, reverse primer for the female-specific splicing variant, and reverse primer for the male-specific variant, respectively (Table S2).

**Table 1.** Results of blast search for OD1 against the genomes and/or transcriptomes of six termite species and their sister group (woodroach).

|  | Database | *dsx* | *Dmrt11* | *Dmrt93* | *Dmrt99* |
| --- | --- | --- | --- | --- | --- |
| *Reticulitermes* | Genome (Rspe OGS1.0) | ***Rspe_dsx*** (scaffold6: | RS007930 | RS006912 | RS002870 |
| *speratus* | transcriptome | 6307974..6309410) |  |  |  |
| *Zootermopsis* | Genome (Znev OGS | No hit | scaffold668 | Znev05388* | Znev16235* |
| *nevadensis* | v2.229) |  |  |  |  |
| *Hodotermopsis* | Transcriptome | ***Hsjo_dsx*** (c35221) | c18070 | c38968 | No hit |
| *sjostedti* |  |  |  |  |  |
| *Nastitermes* | Transcriptome | ***Ntak_dsx*** | comp25194 | comp174542 | No hit |
| *takasagoensis* |  | (G5ZWOJF02FLJ2Z*) |  |  |  |
| *Macrotermes* | Genome | ***Mnat_dsx*** (Mnat_08109) | No hit | Mnat_01812 | Mnat_08410 |
| *natalensis* | (Mnat_gene_v1.2.cds),<br>transcriptome |  |  |  |  |
| *Cryptotermes* | Genome (Csec_1.0), | ***Csec_dsx1*** | XM0238633381 | XM0238639211 | XM0238500541 |
| *secundus* | Transcriptome | (XM_023861307),<br>***Csec_dsx2*** (XM_023858380) |  |  |  |
| *Cryptocercus* | Transcriptome | ***Cpun_dsx*** † | comp1991 | No hit | No hit |
| *punctulatus* |  | (Cpun_comp8195_c0_seq1) |  |  |  |

\* is also described by Price et al. <sup>6</sup>
† found by BLAST search using OD2 (but not OD1) sequences of *Blattella dsx* orthologues, followed by 5' RACE

Phylogenetic analysis of the OD1 nucleotide sequences revealed that all *dsx* reported in holometabolous insects belonged to a single clade (*dsx* clade) (Fig. 2). This clade contained a single sequence containing OD1, but no OD2, from each of the four termites *H. sjostedti* (c35221 ^16^), *R. speratus* (not annotated, scaffold 6 ^20^: 6307974.6309410), *N. takasagoensis* (G5ZWOJF02FLJ2Z ^16^), and *M. natalensis* (Mnat_08109 ^18^); two sequences from *Cryptotermes secundus* (XM_023861307 and XM_023858380^15^) (Fig. 2, Table 1). No sequence from *Z. nevadensis* was detected within this clade. The other OD1 sequences from insects formed clades specific to the *DMRT11*, *DMRT93*, and *DMRT99* orthologues (Fig. 2). To examine whether the *dsx*-clade sequences in termites were *dsx* orthologues, synteny analyses for *dsx* were conducted based on the proximity of the *dsx* and *prospero* (*pros*), which is well conserved among seven different holometabolous and hemimetabolous insect orders ^9^. The gene encoding the *dsx*-clade sequence was located 83 kb and 80 kb close to the *pros* orthologues (RS012493 ^22^ and MN005277 ^23^) in scaffold 6 of *R. speratus* and scaffold 295 of *M. natalensis* (Mnat_08109), respectively ^20^. Therefore, these *dsx-*clade genes were designated as the termite orthologues of *dsx* (*Rspe_dsx* and *Mnat_dsx*, Table 1). In *C. secundus*, both *dsx*-clade sequences, XM_023861307 and XM_023858380, and the *pros* orthologue (XM_023851644) were located in scaffold 829 (length: 88.5 kb), 635 (length: 1.3 Mb), and 511 (length: 2.2 Mb), respectively. Because the scaffold lengths (especially for scaffold 829) might be insufficient for the synteny analysis based on proximity (17–245 kb ^9^), evidence of synteny in *C. secundus* could not be obtained. The synteny in *H. sjostedti* and *N. takasagoensis* could not be examined because of the unavailability of their genome data. On the basis of the OD1 sequence similarity (Fig. 2), we designated these as *dsx* orthologues (hereafter referred to as *Csec_dsx1*, *Csec_dsx2*, *Hsjo_dsx*, and *Ntak_dsx*, Table 1).

**Figure 2.**
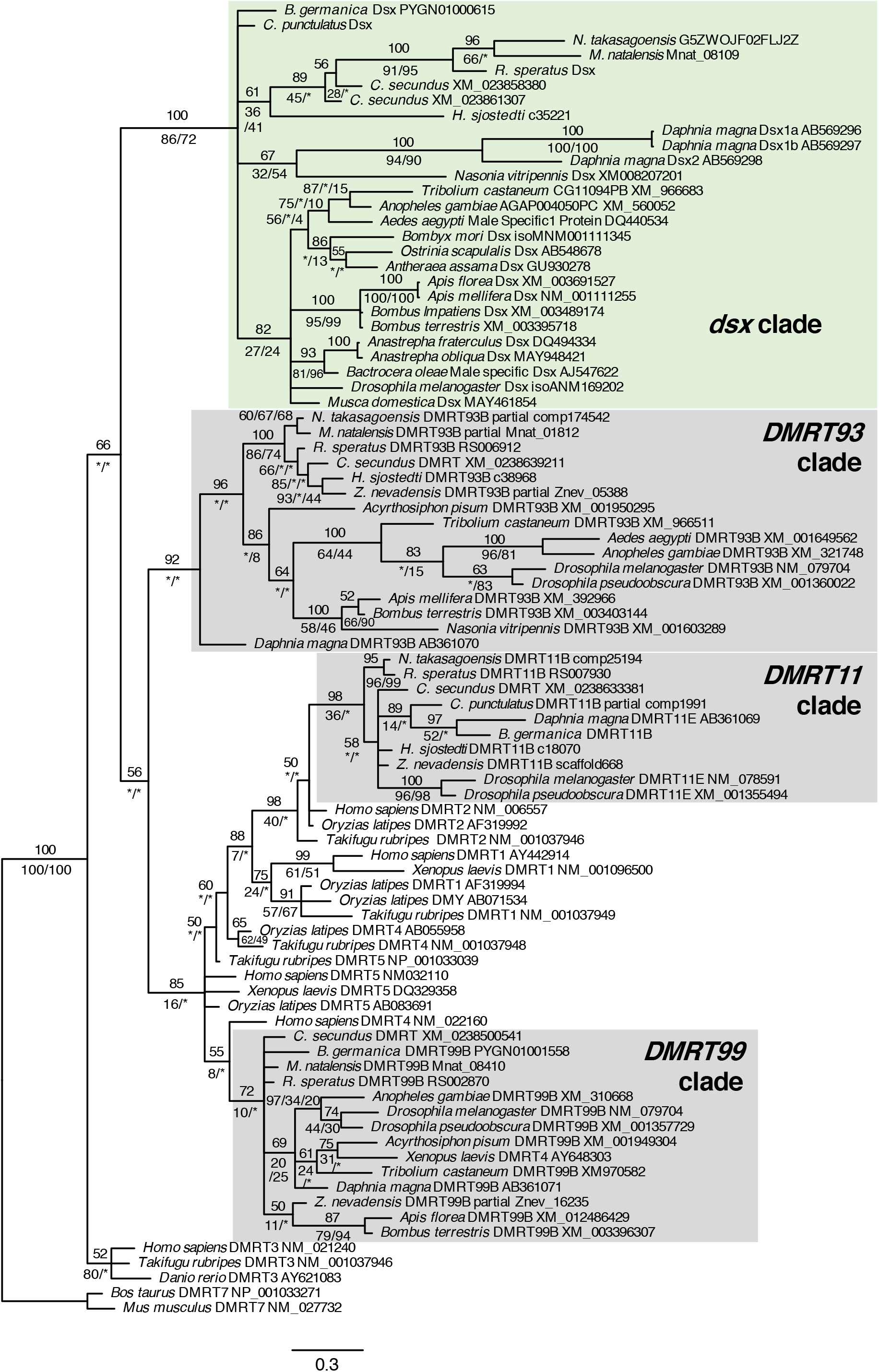
Molecular phylogenetic tree of *dsx* and *DMRT* homologues. Bayesian tree of *dsx* and *DMRT* of insects and crustaceans was constructed based on the OD1 sequences (135 bp with no gaps). Numbers shown above branches represent the Bayes posterior probabilities. Bootstrap values (1,000 replicates) are shown below branches to indicate the level of support in the ML and MP methods, respectively. An asterisk indicates that a node is not supported by the ML and MP methods.

### Splicing patterns and expression of *doublesex* homologues in termites and woodroach

To confirm the sex-specific splicing of *Cpun_dsx*, expression levels of the predicted sex-specific splicing isoforms were quantified by quantitative RT-PCR using primers specific to the predicted isoforms (Fig. 1 and Table S2). The predicted female- and male-specific isoforms was confirmed to be expressed abundantly in females (*t* = −6.68, *p* < 0.01, generalised linear model [GLM]) and males (*t* = 7.88, *p* < 0.01, GLM), respectively (Fig. 3), indicating that *Cpun_dsx* was spliced in a sex-specific manner, as shown in holometabolous insects and the German cockroach *B. germanica* ^9,15^. Next, to obtain the splicing isoforms of *Hsjo_dsx, Rspe_dsx*, and *Ntak_dsx*, we performed 3 ‘-RACE PCRs using gene-specific primers (Table S2) and amplified only the single fragment downstream of OD1 for each species (Fig. 1). The amplified downstream sequences of each species were consistent with those obtained by the aforementioned BLAST search. Although *dsx* is composed of ca. five exons, of which two to three posterior exons are sexually spliced in almost all insects examined previously ^24^, each of the predicted full-length transcripts of *Rspe_dsx, Mnat_dsx, Csex_dsx1*, and *Csec_dsx2* was encoded by a single exon in their genome. Based on the transcriptome data in *R. speratus* ^21^, *Rspe_dsx* was expressed only in male reproductives (primary kings), soldiers, and workers, but not in females, and in both heads (*t* = 3.99, *p* < 0.01, GLM) and the other parts (referred to as just “body”) (*t* = 10.97, *p* < 0.001, GLM) (Fig. 4A). Quantitative RT-PCR also showed male-specific expression patterns even in the nymphs (*t* = 5.69, *p* < 0.01, GLM, Fig. 4B) and eggs (*t* = 3.10, *p* < 0.05, GLM, Fig. 4C). In addition, *Ntak_dsx* was expressed only in males, regardless of caste (*t* = 3.70, *p* < 0.01, GLM, Fig. 4D). These results indicated that termite *dsx* was not alternatively spliced, but transcribed in a male-specific manner.

**Figure 3.**
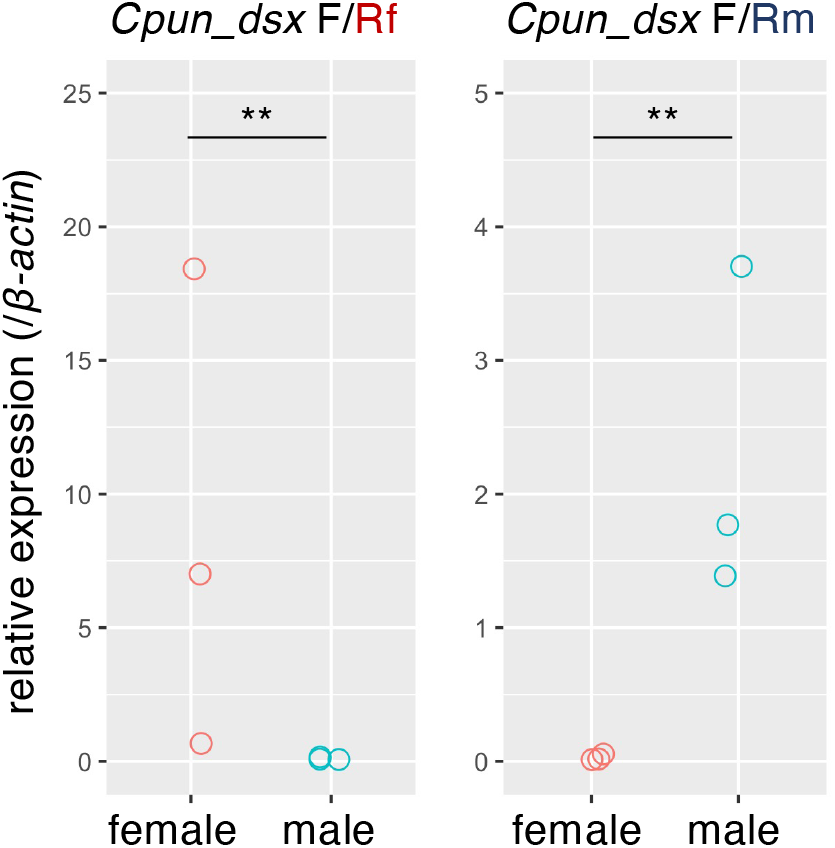
Expression patterns of *Cpun_dsx* in the female and male gonads of *C. punctulatus* adults. Left and right panels show the relative expression levels of its female- and male-specific isoform to those of *β-actin*, respectively. “F”, “Rf”, and “Rm” indicate the primers shown in Fig. 1. The expression levels were compared between sexes using GLM analyses. ** indicates *p* < 0.01.

**Figure 4.**
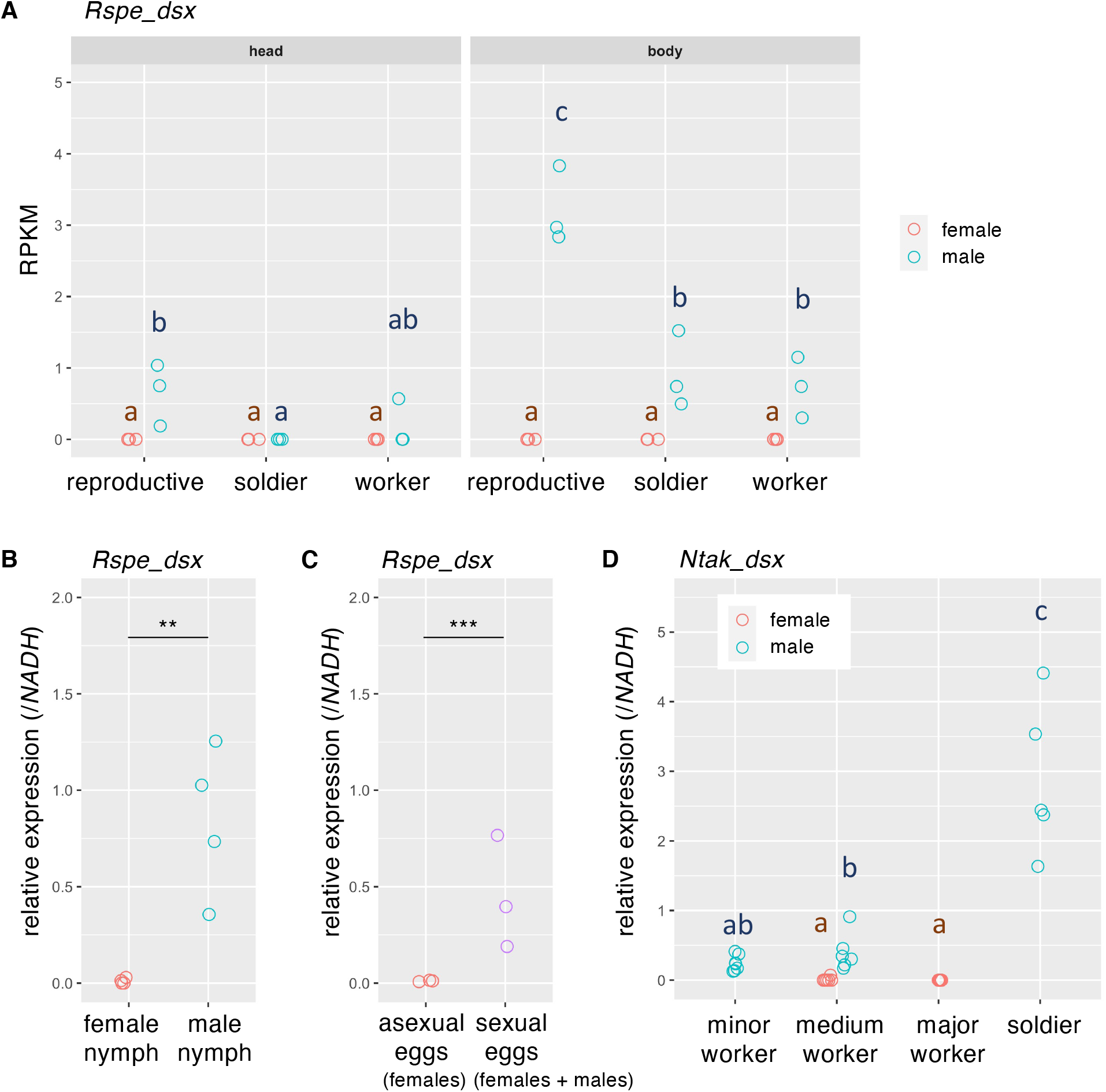
Sex-specific expression patterns of *Rspe_dsx* and *Ntak_dsx*. (A) Reads per kilo base per million mapped reads (RPKM) of *Rspe_dsx* in each sex and in each caste of *R. speratus*, which were re-analysed RNA-seq data (DRA010978, biological triplicates ^21^). (B) *Rspe_dsx* expression levels relative to those of *NADH* in female and male nymphs. (C) Relative *Rspe_dsx* expression in eggs produced asexually and sexually. The formers are fated to develop into females, whereas the sexual fate of the latter was not determined. (D) *Ntak_dsx* expression levels relative to those of *NADH* in each sex and in each caste of *N. takasagoensis*. In (A) and (D), the effects of sex, caste, and their interaction on gene expression levels were evaluated using GLM followed by multiple comparison using linear hypothesis testing with Tukey adjustment in the “multcomp” R package. Different letters indicate significant differences according to multiple comparison (*p* < 0.05). In (B) and (C), relative expression levels of *Rspe_dsx* were compared between sexes using GLM analyses. ** and *** indicates *p* < 0.01 and 0.001 in GLM, respectively.

### Predicted binding sites and expression patterns of putative regulatory factors for male-specific transcription of *Rspe_dsx*

To examine the proximate mechanism for male-specific transcription of termite *dsx*, we searched for putative transcription factors that bind to the promoter region of *Rspe_dsx*, and those expressed in a male- or female-specific manner. First, 1 kb upstream of the transcription start site of *Rspe_dsx* was extracted as a transcriptional regulatory region, as defined by Toyota et al. ^25^. Next, *de novo* motif discovery was performed for the promoter using hypergeometric optimisation of motif enrichment (HOMER) v4.11 ^26^, based on the equipped motif library of known transcription factors in insects. Although 32 putative TFBSs were detected in the promoter, 20 homologues out of the 32 transcription factors existed in the *Reticulitermes* genome, of which five were duplicated in the genome (Table 2). Based on the transcriptome data in *R. speratus* ^21^, however, none of them were expressed in a sex-specific manner in both heads and bodies, except for the *vielfaltig* (*vfl*) orthologue (GLM, Table 2). The expression levels of the *vfl* orthologue were significantly different between sexes only in the bodies of reproductives, but similar patterns were not observed in other castes (Fig. S1).

**Table 2.**
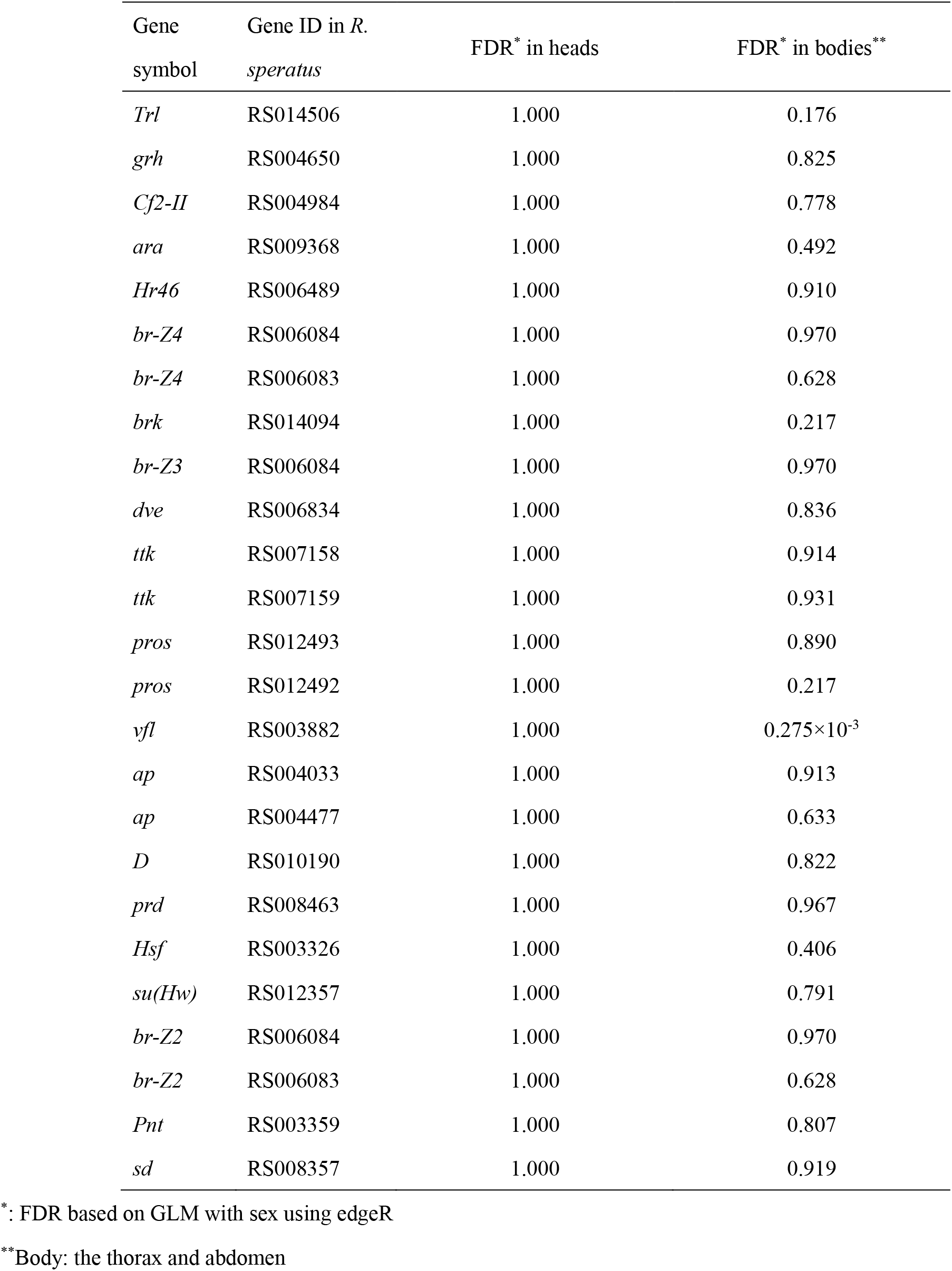
Transcription factors with putative binding sites detected in the *R. speratus* genome and their expression differences between sexes.

## Discussion

The *dsx* orthologue of the woodroach was spliced in a sex-specific manner, similar to that of the German cockroach and holometabolous insects (Fig. 5). Surprisingly, however, termite *dsx* homologues did not exhibit sex-specific splicing, and their transcription was regulated in a male-specific manner. Male- or female-biased expression of *dsx* has been reported in some arthropods, for example, water fleas ^5,25^, red claw crayfish ^27^, Chinese shrimp ^28^ (Crustacea), and mites ^7^ (Chelicerata), suggesting that sex-biased *dsx* transcriptions are an ancestral state in Arthropoda ^9^. In contrast, *dsx* produces sex-specific splice isoforms in most hemimetabolous and holometabolous insects examined to date ^6,9,24^ except for the lice *Pediculus humanus* and the hemipteran *Bemisia tabaci*. These two hemimetabolous insects have *dsx* homologues with non-sex-specific isoforms ^8,9^. Consequently, the sex-specific alternative splicing of *dsx* would be acquired early in the evolution of insects, and secondarily lost in some hemimetabolous insects ^9^, including termites. Within the cockroach and termite clade, after the ancestral termites had derived from the common ancestor of *Cryptocercus* and termites (around 140.6 MYA, ranging from 112.6 to 170.5 MYA ^29^), it is suggested that the sex-specific *dsx* splicing was lost, and the male-specific *dsx* transcription was acquired alternatively in termites (Fig. 5). No *dsx* orthologues in *Z. nevadensis* are mystery at present, and further exhaustive search should be needed.

**Figure 5.**
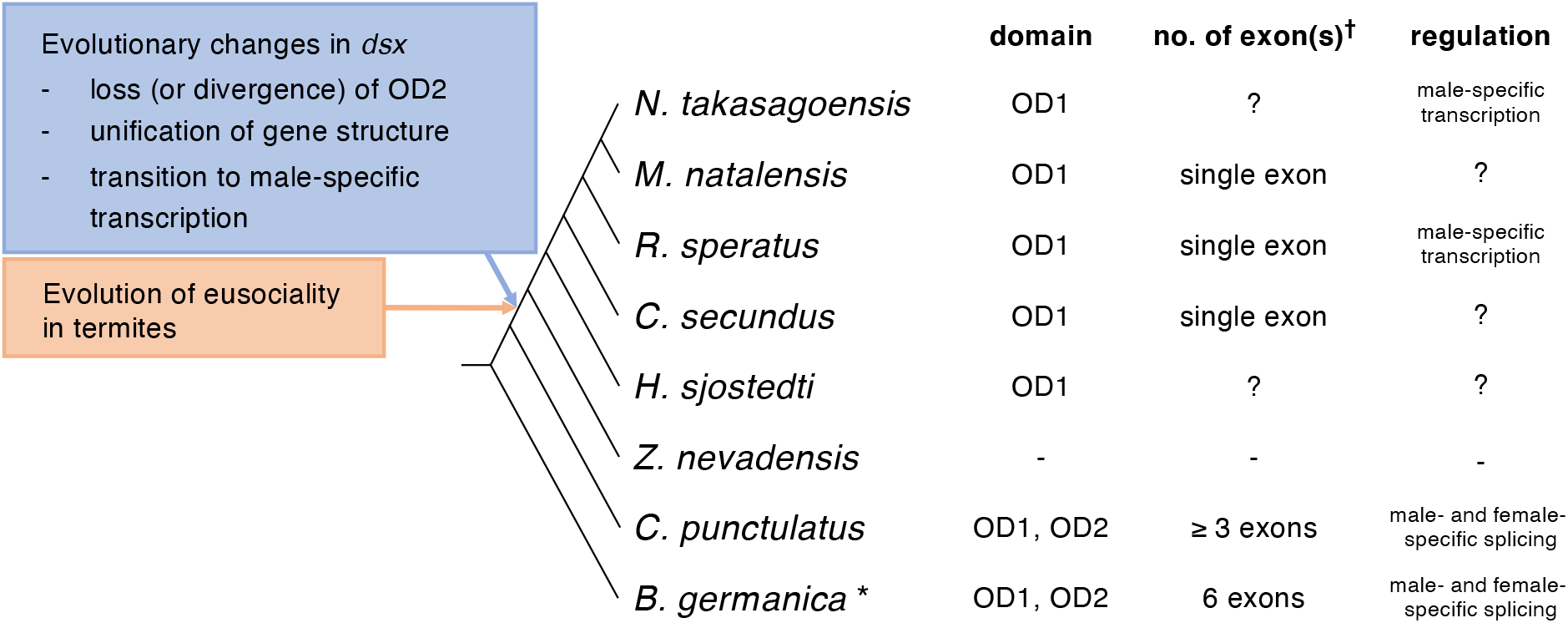
Evolutionary changes in *dsx* regulations associated with social evolution in termites. The presence of OD1 and OD2, numbers of exons containing coding sequences (†), and regulatory manners of *dsx* homologues were compared among German cockroach, woodroach, and six termite species. “-” and “?” mean “not detected” and “unknown”, respectively. *: Data on *B. germanica* were cited from Wexler et al. ^9^. The tree topology was based on that of Bourguignon et al. ^52^ and Bucek et al. ^29^.

Although *dsx* consists of approximately five exons in most insects ^24^, two *dsx* paralogs (*dsx1* and *dsx2*) in water fleas consist of four and two exons, respectively ^5^. Its homologue in Chinese shrimp consists of two exons ^28^, and those in mites ^7^ and termites consist of only a single exon (Fig. 5). Moreover, the present study showed no homologous OD2 in termite *dsx* (Fig. 5), suggesting that OD2 has indeed been lost or has diverged beyond our ability to detect it ^6^. OD2 is conserved in *dsx* orthologues with sex-specific splicing in holometabolous insects, as well as in those with sex-specific transcription in water fleas ^5,25^ and Chinese shrimp ^28^ but not in red claw crayfish ^27^. Both OD1 and OD2 are related to bind target DNA as a dimeric DNA binding unit in the fruit fly *D. melanogaster*: OD1 enables Dsx to bind target DNA and dimerises weakly in a DNA-dependent manner ^12^, whereas OD2 is necessary for dimerisation and enhances DNA recognition of OD1 ^30^. In addition, fly OD2 contains sex-specific spliced sequences, which may be involved in the formation of sex-specific units for transcriptional regulation of downstream target genes through either its sex-specific interactions with the transcriptional machinery or its sex-specific DNA binding ^12^. The decreasing numbers of *dsx* exons in some arthropods and termites and the loss (or divergence) of OD2 in termites would be associated with the evolution of *dsx* regulation, although it remains unknown whether such evolutionary changes in the gene structure were the cause or consequence of the loss of sex-specific splicing.

The sex determination cascade has never been examined in termites. Almost all orthologues of sex determination genes reported in holometabolous insects (e.g., *transformer*, *transformer-2*, and *Sex-lethal*) are conserved in the genome of *R. speratus* ^22^. However, these orthologues encode splicing factors and are unlikely to directly regulate male-specific *dsx* transcription. Male-specific transcription should be regulated by a transcription factor expressed in a sex-specific manner in every caste, as shown in termite *dsx*. Although we found 27 putative TFBSs in the *Rspe_dsx* promoter region, no orthologues of 27 transcription factors showed sex-specific expression patterns similar to those of *Rspe_dsx*. However, our motif search was based on the binding motifs found in the fruit fly *D. melanogaster*. In addition, unidentified binding motifs could be present out of the transcriptional regulatory region examined in this study. Further investigations are needed to determine whether the unidentified transcription factors expressed in a sex-specific manner would regulate male-specific *dsx* transcription in termites.

The transition of *dsx* regulation in termites is associated with the social evolution of cockroaches and termites (Fig. 5). Cooperative brood care by parents and generation overlap should have already been acquired in the last common ancestor of *Cryptocercus* and termites ^31^. Therefore, the male-specific transcription of termite *dsx* would play a role in the regulation of reproductive division of labour, especially the fortress defence, with an altruistic sterile “soldier” caste that had occurred at the origin of Isoptera ^32^. Some species, especially in the most derived family (Termitidae), have sex-specific or sex-biased soldier ratios and soldier differentiation pathways ^33^. For example, all soldiers are females in most species of Termitinae and Macrotermitinae, whereas all soldiers are males in most of the examined species in Nasutitermitinae. Soldier differentiation requires high juvenile hormone titres in workers, and the strongly biased soldier-sex ratio might be caused by the differences in juvenile hormone titres (and probably related gene expression levels) between male and female workers ^34,35^. In addition, reproductive caste differentiation is regulated in a sex-specific manner (Oguchi et al. ^36^). Some “queen genes” with high expression levels in female reproductives are identified in some species (reviewed in Korb ^37^). Furthermore, the upregulated genes in male reproductives, which are probably involved in male fertility, are also identified in species with available genome sequences ^15,19,22^. Termite *dsx* could regulate this sexually dimorphic expression by upregulating or downregulating the expression of its target genes in a male-specific manner, resulting in reproductive and non-reproductive division of labour.

## Materials and Methods

### Insects

Seven mature colonies of *R. speratus* were collected in Toyama Prefecture in 2016 and 2019. One *H. sjostedti* mature colony was collected in Yakushima Island, Kagoshima Prefecture in 2015. Two *N. takasagoensis* mature colonies were found on Ishigakijima Island, Okinawa Prefecture, in 2017 and 2018. They were brought back to the laboratory and kept at ca. 25°C in constant darkness. Three families of *C. punctulatus* were collected at Mountain Lake Biological Station, Giles County, VA in April 2015-2017 ^38^ and kept at 15°C in constant darkness. Testes and ovaries were dissected from three males and three females, respectively, and stored at −80°C until RNA extraction for subcloning and expression analyses of *Cpun_dsx*.

### *Blast searches for* dsx *and* dmrt *homologues in termites and cockroaches*

tBlastX searches were performed against genome and/or transcriptome databases of six termite species and one woodroach species (Table 1), using SequenceServer ^39^. Either the OD1 (45 amino acids) or OD2 (45 amino acids) sequences of *B. germanica dsx* were used as queries. Sequences with an E value less than 10^-5^ were selected as candidates for *dsx* orthologues and used for phylogenetic analysis.

### *Construction of phylogenetic tree of OD1 sequences and identification of* dsx *orthologues*

To identify the *dsx* orthologue, a phylogenetic tree of OD1 sequences (135 bp with no gaps) was constructed according to previous studies ^5,40,41^. We used 84 OD1-containing genes (Table S3 ‘OTU’). The DMRT7 genes of vertebrates (mouse and cattle) were used as the outgroups. Phylogenetic relationships were inferred using Bayesian inference (BI), maximum likelihood (ML), and maximum parsimony (MP) methods. For BI, the most appropriate model of sequence evolution was determined using the model selection option implemented in MEGA version 7.0.21 ^42^, and the GTR + G model was selected. Using MrBayes version 3.2.6 ^43^, a total of 100,000 trees were obtained (ngen = 10,000,000, samplefreq = 100). The first 25% of these (25,000) were discarded as burn-ins, and a 50%-majority-rule consensus tree was produced. For ML, 1,000 bootstrap replicates were performed based on the same model of sequence evolution as BI in MEGA 7.0.21, with the default tree inference options. For MP analysis, all characters were included and weighted equally, and 1,000 bootstrap replicates were performed using MEGA 7.0.21. The Subtree-Pruning-Regrafting algorithm, in which the initial trees were obtained by the random addition of sequences (10 replicates), was used.

### *Subcloning of the candidate* dsx *orthologues*

Female and male alate reproductives (winged adults) of *R. speratus* were obtained from three colonies collected in 2016. According to previous studies ^44,45^, incipient colonies were established using unrelated female and male alates. Total 30 eggs and primary reproductives (queen and king) were obtained from 20 colonies at 1.5 months after colony establishment. Total RNA of the eggs was extracted and treated with DNase I using the ReliaPrep RNA Tissue Miniprep System (Promega, USA). Total RNA of the primary queens or kings was separately extracted from the body (except for the head parts) of three individuals using ISOGEN II (Nippongene, Japan). Total RNA of female or male alates of *N. takasagoensis* (using the colony collected in 2017) was separately extracted from the body (except for the head parts) of five individuals using ISOGEN II. Total RNA from the secondary queen or king of *H. sjostedti* (using the colony collected in 2015) was separately extracted from the body (except for the head part) of one individual using ISOGEN II. Female and male *C. punctulatus* adults were dissected, and the gonads (ovaries or testes) were collected from each individual. Total RNA was extracted from the gonads using ISOGEN I and II, and then treated with RNase-free DNase I (Takara, Japan). The purity and quantity of the extracted RNA were measured using a NanoVue spectrophotometer (GE Healthcare Bio-Sciences, Japan).

The 5’ and 3’ ends of *Cpun_dsx* were amplified using the SMARTer RACE 5’/3’kit (TaKaRa, Shiga, Japan) and gene-specific primers designed for OD2 (*“Cpun_dsx* OD2 5’RACE” and *“Cpun_dsx* OD2 3’RACE”, Table S1). Its male-specific exon was amplified using Advantage^®^ 2 Polymerase Mix (TaKaRa), the gene-specific primer *“Cpun_dsx* OD2 3’RACE”, and a reverse primer in the male-specific exon of *Blattella dsx*, which was located at terminal codon and 3’UTR (“*Cpun*_*dsx* male-specific exon-R”, Table S1). The 3’ ends of *Rspe_dsx, Hsjo_dsx*, and *Ntak_dsx* were also amplified using the SMARTer RACE 5’/3’ Kit and newly designed gene-specific primers (Table S1). The amplified fragments were subcloned into the pGEM-T easy vector system (Promega), and the nucleotide sequences of each fragment were determined using ABI Prism Big Dye Terminator v3.1 Cycle Sequencing Kit in conjunction with a 3500 Genetic Analyzer (Applied Biosystems, Foster City, CA, USA). The newly identified gene sequences were deposited in the GenBank/EMBL/DDBJ database (Table 1).

### RNA-seq analysis

RNA-seq data of *R. speratus* were used to compare the expression levels of *Rspe_dsx* (Table 1) among three castes (workers, soldiers, and reproductives), two body parts (heads and the remaining parts), and two sexes (biological triplicates; NCBI BioProject Accession No. PRJDB5589). The filtered RNA-seq reads were mapped onto their genome assembly ^20^ using TopHat v2.1.021. Transcript abundances were then estimated using the featureCounts program of the Subread package (Liao et al. 2013). To compare gene expression levels among castes and between sexes, first, counts per million (CPM) were calculated from the estimated transcript abundances. Genes with at least CPM of 1 in at least three samples were kept for subsequent analyses. CPM values were then normalized with the trimmed mean of M-values (TMM) algorism in edgeR (Robinson et al. 2010). Differential gene expression analyses were performed separately for each body parts using a GLM with two factors, namely, caste and sex implemented in edgeR, and then, genes with false discovery rate (FDR) < 0.01 were identified as genes expressed in a sex-specific manner. RPKM (Reads Per Kilobase Million) values were calculated by dividing the CPM values by the length of the genes in kilobases.

### Real-time quantitative PCR

Gene-specific primers were designed against *Rspe_dsx, Ntak_dsx*, and *Cpun_dsx* using Primer3Plus ^46^ for real-time quantitative PCR (Table S2). Total RNA of female and male nymphs of *R. speratus* (from the colony collected in 2019) was separately extracted from the whole body of 10 individuals using ISOGEN II. Total RNA of male (minor and medium) workers, female (medium and major) workers, and male soldiers of *N. takasagoensis* (from the colony collected in 2018) were separately extracted from the whole body of five individuals using ISOGEN II. Total RNAs derived from the gonads of adult woodroaches were also used. DNase treatment was performed using the same method described above. According to previous studies ^44,45^, incipient colonies (queen-king and two-queen colonies) of *R. speratus* were established using alates derived from three colonies collected in 2019. Total RNA of eggs produced asexually and sexually, which develop into females and both sexes, respectively, were extracted from 30 eggs and treated with DNase I using ReliaPrep RNA Tissue Miniprep System (Promega). cDNAs were synthesised from the purified RNA using a High-Capacity cDNA Reverse Transcription Kit (Applied Biosystems). Biological replications (n = 3-5) were prepared for each category. Expression analyses were performed using Thunderbird SYBR qPCR Mix (Toyobo, Japan) with a Mini Option Real-time PCR system (Bio-Rad, Japan) and an Applied Biosystems 7500 Fast Real-Time PCR System (Applied Biosystems).

To determine a sustainable internal control gene for *R. speratus*, the expression levels of six genes *[EF1-α* (accession no. AB602838), *NADH-dh* (no. AB602837), *β-actin* (no. AB520714), *GstD1* (gene id RS001168), *EIF-1* (RS005199), and *RPS18* (RS015150)] were evaluated using the GeNorm ^47^ and NormFinder ^48^ software (Table S4). For *C. punctulatus* and *N. takasagoensis*, according to previous studies ^38,49^, three genes [*β-actin* (nos. Cp_TR19468 and AB501107), *NADH-dh* (Cp_TR49774 and AB50119), and *EF1-α* (AFK49795 and AB501108)] were evaluated (Tables S5 and S6). All gene-specific primers were designed using Primer3Plus (Table S2). We confirmed the amplification of a single PCR product using dissociation curves and product sizes.

The expression levels were statistically analysed using GLMs with gamma error distribution and log link function. The fixed effects were sex, caste, and their interaction. Multiple comparisons were performed using general linear hypothesis testing (glht function) with Tukey adjustment in the “multcomp” R package ^50^. These analyses were conducted using R ver. 4.0.3 (available at http://cran.r-project.org/).

### Search for transcription factor binding motif

The *Reticulitermes* genome was loaded into the database of HOMER v4.11 using the “loagGenome.pl” program. The “findMotifsGenome.pl” program was executed to search for *Drosophila* motif collections using the “-mcheck” option against the promoter of *Rspe_dsx*, which was the 1.0 kbp upstream region from its transcription start site. The nucleotide sequences of *Drosophila* transcription factors that were detected in their binding sites were subjected to a BlastX search for orthologues against the protein database of *R. speratus* ^38^ using BLAST+ (2.10.1) ^51^. The expression levels of these orthologues were extracted from their transcriptome data ^27^ and statistically analysed using GLMs with gamma error distribution and log link function, as mentioned above.

## Acknowledgement

Chrisnine A. Nalepa kindly provided us with *Cryptocercus punctulatus*. We thank Shutaro Hanmoto for his assistance with laboratory work. We also thank Toru Miura and Masatoshi Matsunami for their constructive suggestions regarding this study. This study was partly supported by JSPS KAKENHI (Grant Numbers JP19K06860 to SM, JP20K06816 to YH, and JP19H03273 to KM) and by the NIBB Collaborative Research Program (Nos. 19-335 and 20-323).

SM, HG, and KM planned and designed the experiments.

## Author contributions

SM, KF, KK, and YM collected the samples. SM, KF, KK, and YM performed the experiments. SM analysed the data. SM, KF, and KM wrote the manuscript. All authors read and approved the final manuscript.

## Another Information

All authors declare that they have no conflict of interest.

---

**Figure S1.**
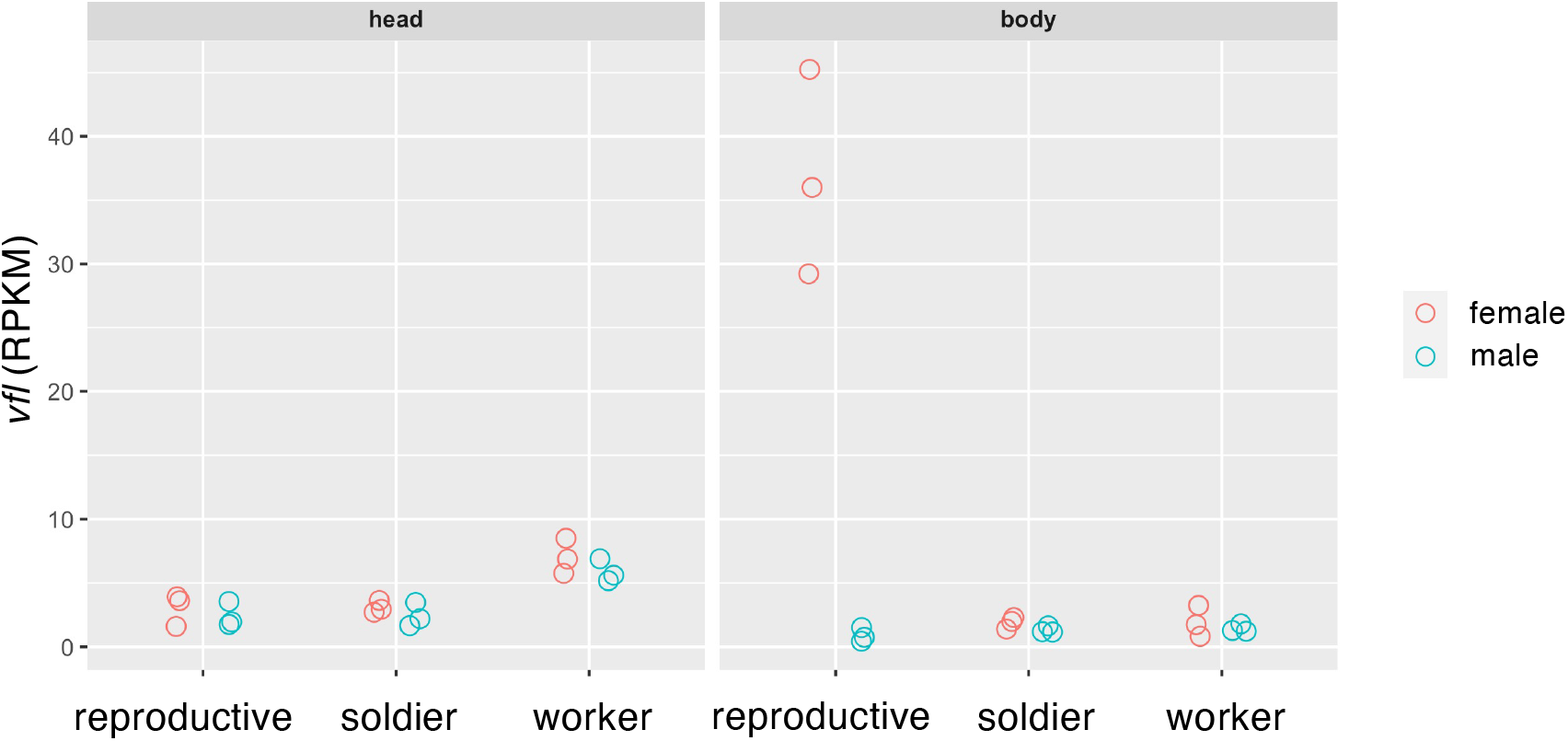
RPKM of *vfl* orthologue in each sex and in each caste of *R. speratus*, which RNA-seq data were deposited in NCBI (DRA010978, biological triplicates ^21^). The effects of sex, caste, and their interaction on gene expression levels were evaluated using GLM.

**Table S1.**
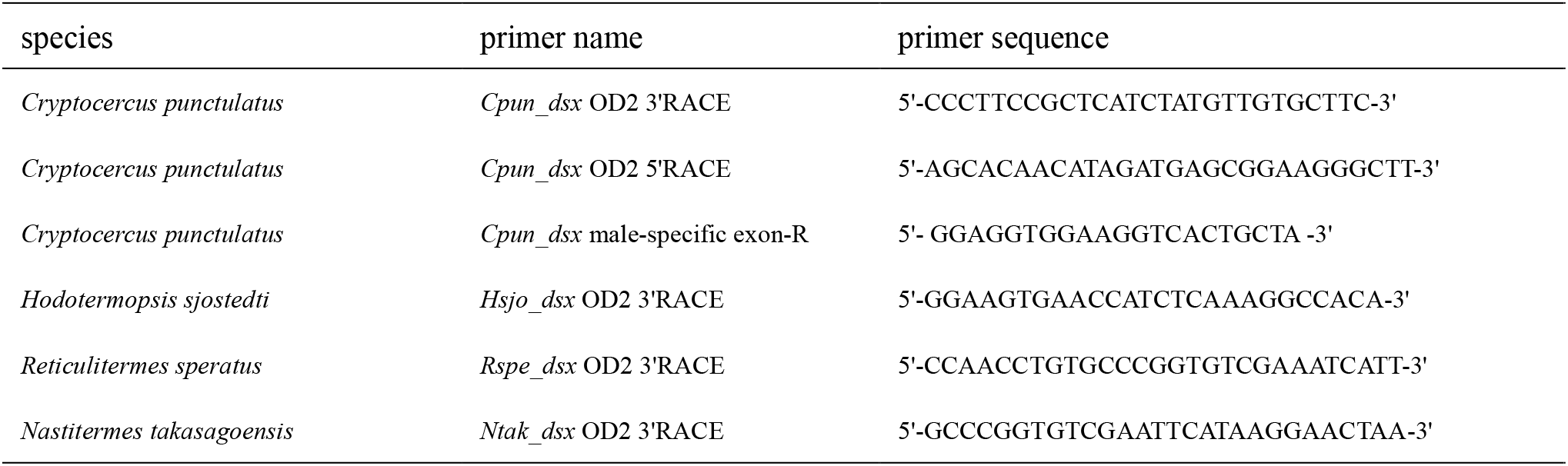
Primers used for RACE PCRs and subcloning.

**Table S2.**
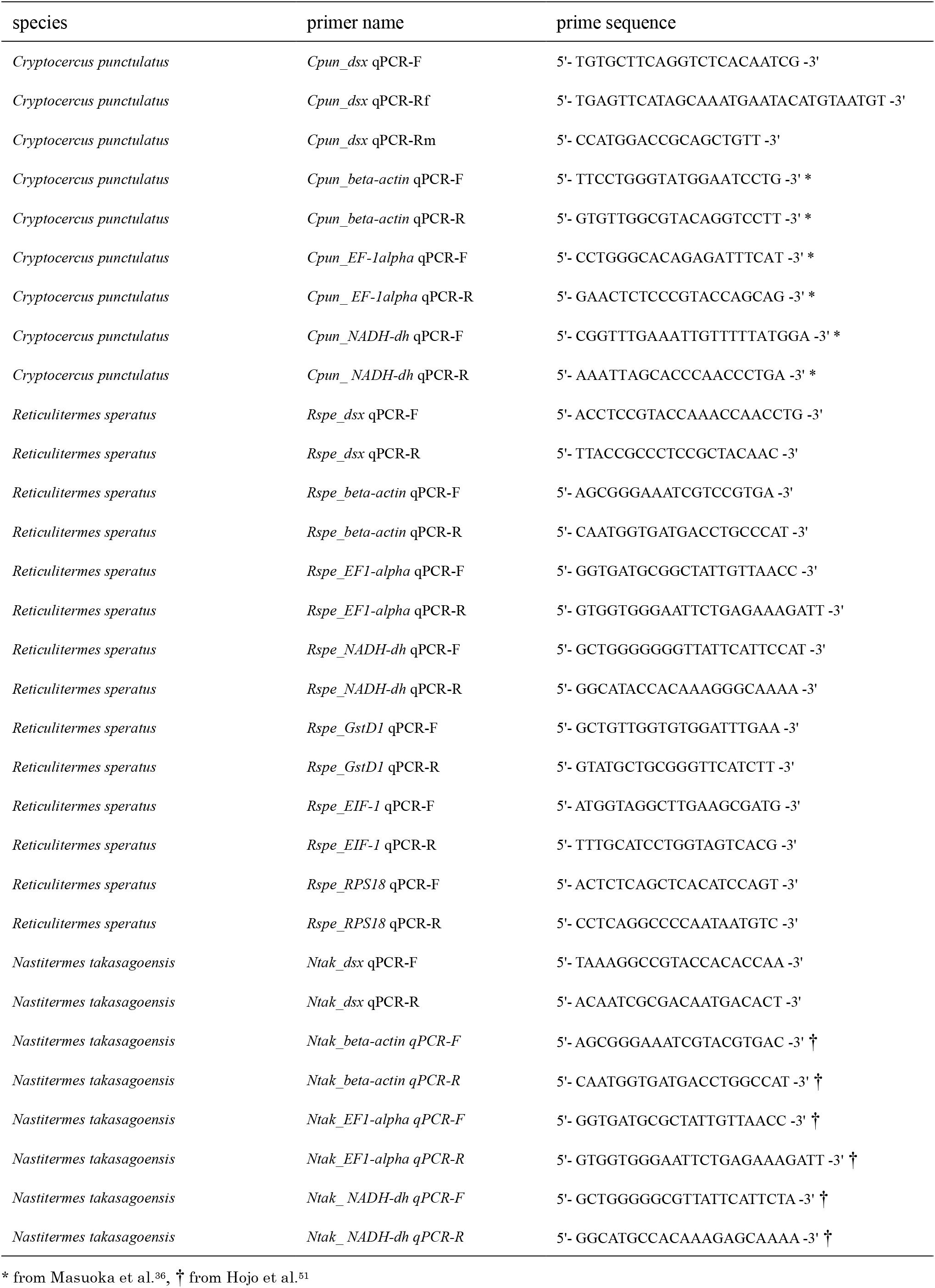
Primers used for quantitative RT-PCRs.

**Table S3.**
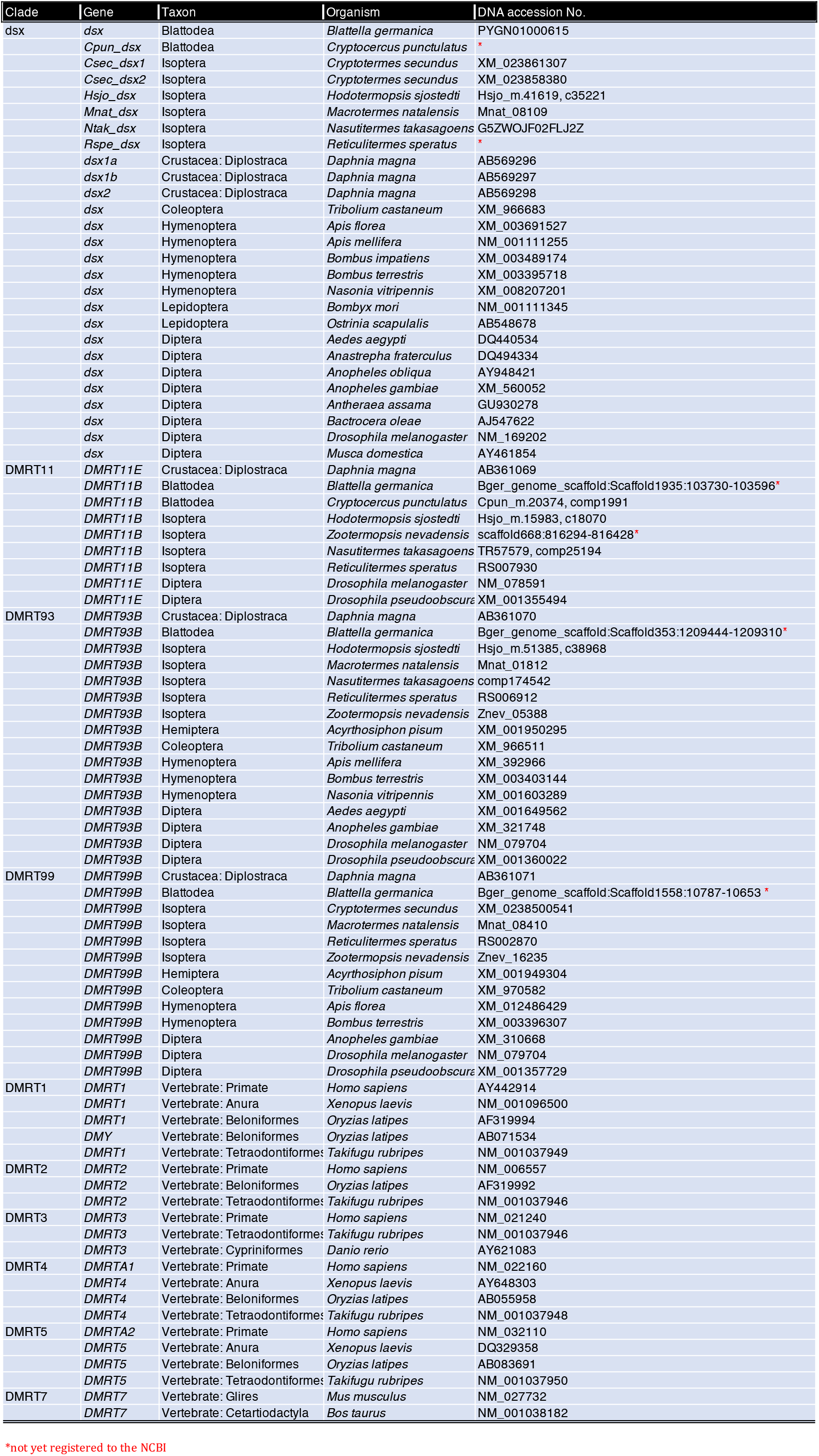
Information of OTU used for phylogenetic analysis.

**Table S4.** Stability values of internal control gene candidates of *R. speratus* using GeNorm and NormFinder.

|  | GeNorm (stability value) | NormFinder (stability value) |
| --- | --- | --- |
| *Rspe_beta-actin* | 0.385 | 0.264 |
| *Rspe_EF1-alpha* | 0.246 | 0.106 |
| *Rspe_NADH-dh\** | 0.148 | 0.051 |
| *Rspe_GstD1* | 0.158 | 0.089 |
| *Rspe EIF-1* | 0.148 | 0.061 |
| *Rspe_RPS18* | 0.161 | 0.093 |

\* *Rspe\_NADH-dh* was selected by GeNorm and NormFinder due to the lowest stability values.

**Table S5.**
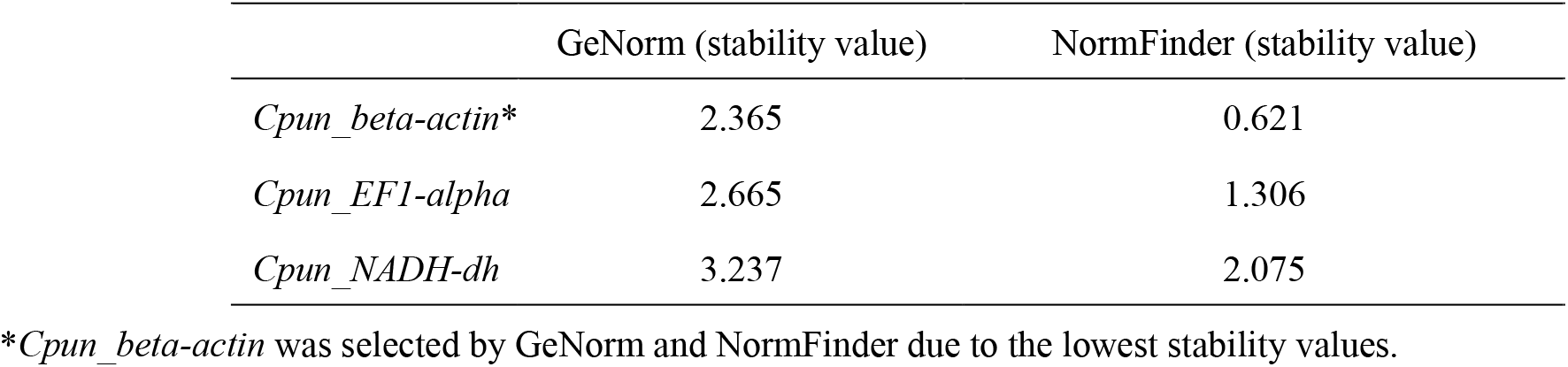
Stability values of internal control gene candidates of *C. punctulatus* using GeNorm and NormFinder.

**Table S6.**
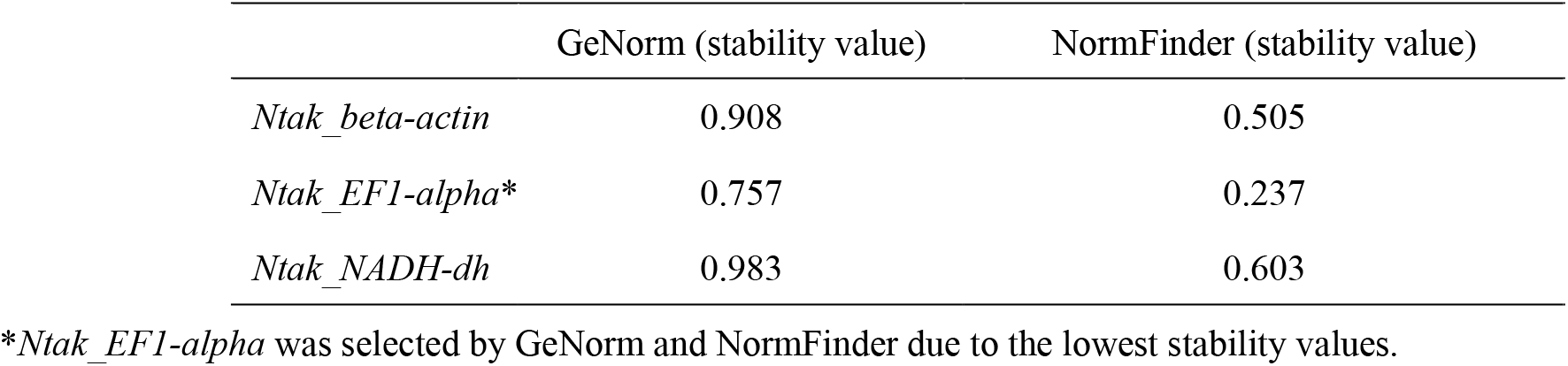
Stability values of internal control gene candidates of *N. takasagoensis* using GeNorm and NormFinder.

